# EXCAVATE-HT: A Bioinformatic Pipeline to Identify Targetable Genomic Variants for Allele-Specific Editing

**DOI:** 10.64898/2026.07.23.739930

**Authors:** Akshita G Saxena, Grace D Ramey, John A Capra, Bruce R Conklin, Bria L Macklin

## Abstract

Allele-specific CRISPR/Cas editing is a powerful tool with great potential for treating genetic diseases and for uncovering the effects of allelic diversity. By targeting commonly inherited single nucleotide polymorphisms (SNPs), a small number of gRNAs can treat many more individuals than targeting rare disease mutations. However, current tools for identifying common targetable variants and generating CRISPR guide RNAs (gRNA) have fundamental conceptual and technical limitations. Here, we introduce EXCAVATE-HT (<u>EX</u>tracting <u>C</u>ommon <u>A</u>llelic <u>VA</u>riants for <u>T</u>argeted <u>E</u>diting in <u>H</u>igh-<u>T</u>hroughput) a bioinformatic tool that mines population variant data to generate CRISPR libraries targeting genomic loci for allele-specific editing. Users define their loci of interest, Cas species, and SNP frequency, then EXCAVATE-HT outputs an annotated list of allele-specific gRNAs. EXCAVATE-HT can also generate libraries of gRNA pairs to enable excision. We illustrate the use of EXCAVATE-HT to design and characterize multiple gRNA libraries for allele-specific targeting of the disease gene, Cone-Rod Homeobox (*CRX)*. EXCAVATE-HT revealed multiple excisions that could treat >30-fold more patients than targeting a single CRX disease mutation.

## INTRODUCTION

Diversity in the human genome has been largely considered a potential complication for genome editing^1,2^. We argue instead that genomic diversity is an asset that can be utilized to enable precise allele-specific editing. Large population genetic studies that recruit diverse participants enable us to identify commonly inherited SNPs across the entire genome and across multiple ancestries. Tens of millions of common SNPs are segregating in human populations, and each individual human genome is heterozygous at between 1.5 and 3 million sites^4^. Moreover, as the cost of whole-genome sequencing continues to decrease, identifying SNPs for personalized therapeutic editing is likely to become routine. Thus, genome editing that targets SNPs offers the opportunity to greatly expand the landscape of gene therapies.

Although the opportunity is great, creating CRISPR therapies that leverage common genomic variation can be daunting, as it requires large-scale and unbiased testing of variant-targeted gRNAs. Screening candidate gRNAs for gene therapies with the greatest clinical impact requires: (1) identifying commonly variable genetic loci from population data, (2) verifying their targetability by CRISPR-Cas systems, (3) identifying the heterozygous common variants present in the genomes of experimental model systems, and (4) computing off-target risk in a high-throughput manner. These processes are challenging without the right software tools and arduous to perform manually.

A limited number of software tools are currently available for designing CRISPR gRNAs that target genetic variation^5–8^ (**Table 1**). These tools enable the generation of a gRNA at a specific locus or the generation of gRNAs targeted to specific SNPs and compatible with multiple Cas species. However, they often limit the number of SNP targets they can take as input or the length of the genomic locus of interest. These tools are also particularly poorly suited for dual-nuclease excision strategies, as they do not allow the pairing of gRNAs in a random or strategic manner. No single tool allows for the mining of both population and personalized variants according to minor allele frequency (MAF), accommodates different Cas species, gRNA length and seed region, and outputs optimized single gRNAs or paired gRNAs, all in a high-throughput manner.

**Table 1.** Capabilities of EXCAVATE-HT and existing software.

| Software Package | SNP Input | Maximum # of SNP inputs | MAF Input | Cas Species Selection | gRNA Library Output | gRNA Pairing | Notes |
| --- | --- | --- | --- | --- | --- | --- | --- |
| CriSNPr<sup>7</sup> | Yes | 1 | Yes | Yes | No | No |  |
| SNP-CRISPR<sup>5</sup> | Yes | 2000 | No | No | No | No |  |
| CRISPAM<sup>8</sup> | Yes |  | No | No | No | No | Only identifies SNPs in PAM |
| CRISPR-HAWK<sup>6</sup> | Yes | No Limit | Yes | Yes | Yes | No |  |
| EXCAVATE-HT | Yes | No Limit | Yes | Yes | Yes | Yes |  |

To address this need, we developed EXCAVATE-HT, an open-source command-line tool for the rapid design of allele-specific CRISPR gRNA libraries. For a given genome, locus of interest, population and/or personalized variant data, and a user-input allele frequency threshold, EXCAVATE-HT generates single and paired gRNA libraries targeting genetic variation (**Figure 1, Figure 2A, B**). The user can select the CRISPR-Cas enzyme as one of four established Cas species (SpCas9, SpCas9 NG, SaCas9, or enAsCas12a) or provide custom PAM sequences and their orientations. At the same time, the user can also input gRNA length and seed region.

**Figure 1.**
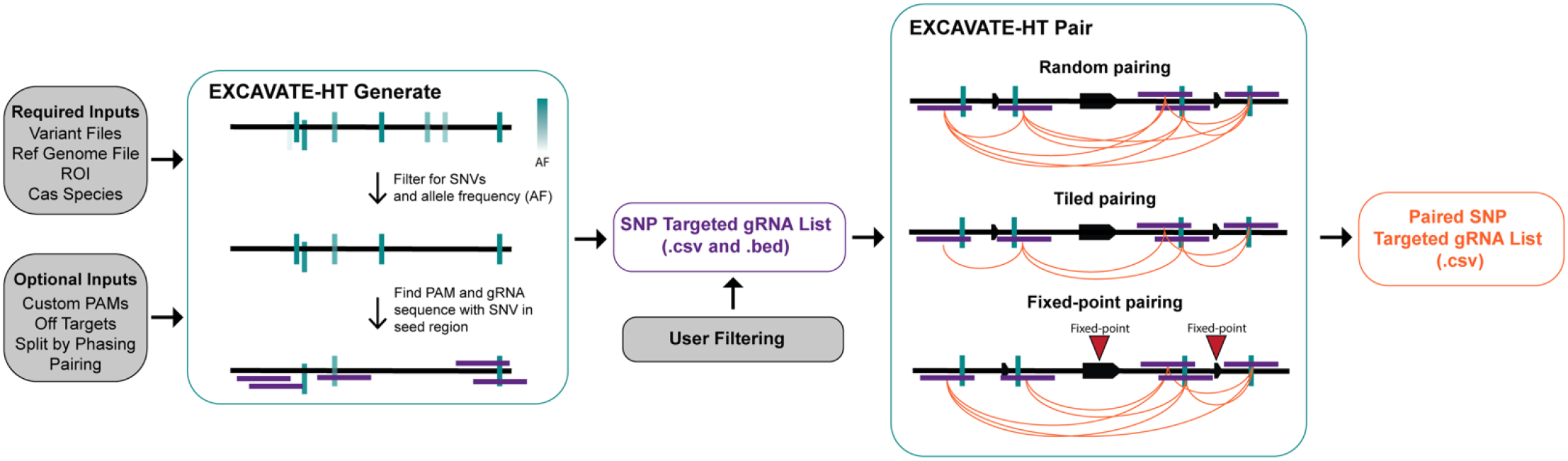
Schematic Outlining EXCAVATE-HT workflow. Required inputs (variant files, reference genome, ROI, and Cas species) are processed by EXCAVATE-HT Generate, which filters variants for SNVs by minor allele frequency and identifies gRNA sequences harboring the SNV in the seed region or PAM. The SNP-targeted gRNA list is exported as .csv and .bed files and optionally filtered by the user before pairing. EXCAVATE-HT Pair then generates paired gRNA libraries using random, tiled, or fixed-point pairing strategies, outputting a paired SNP-targeted gRNA list (.csv).

**Figure 2.**
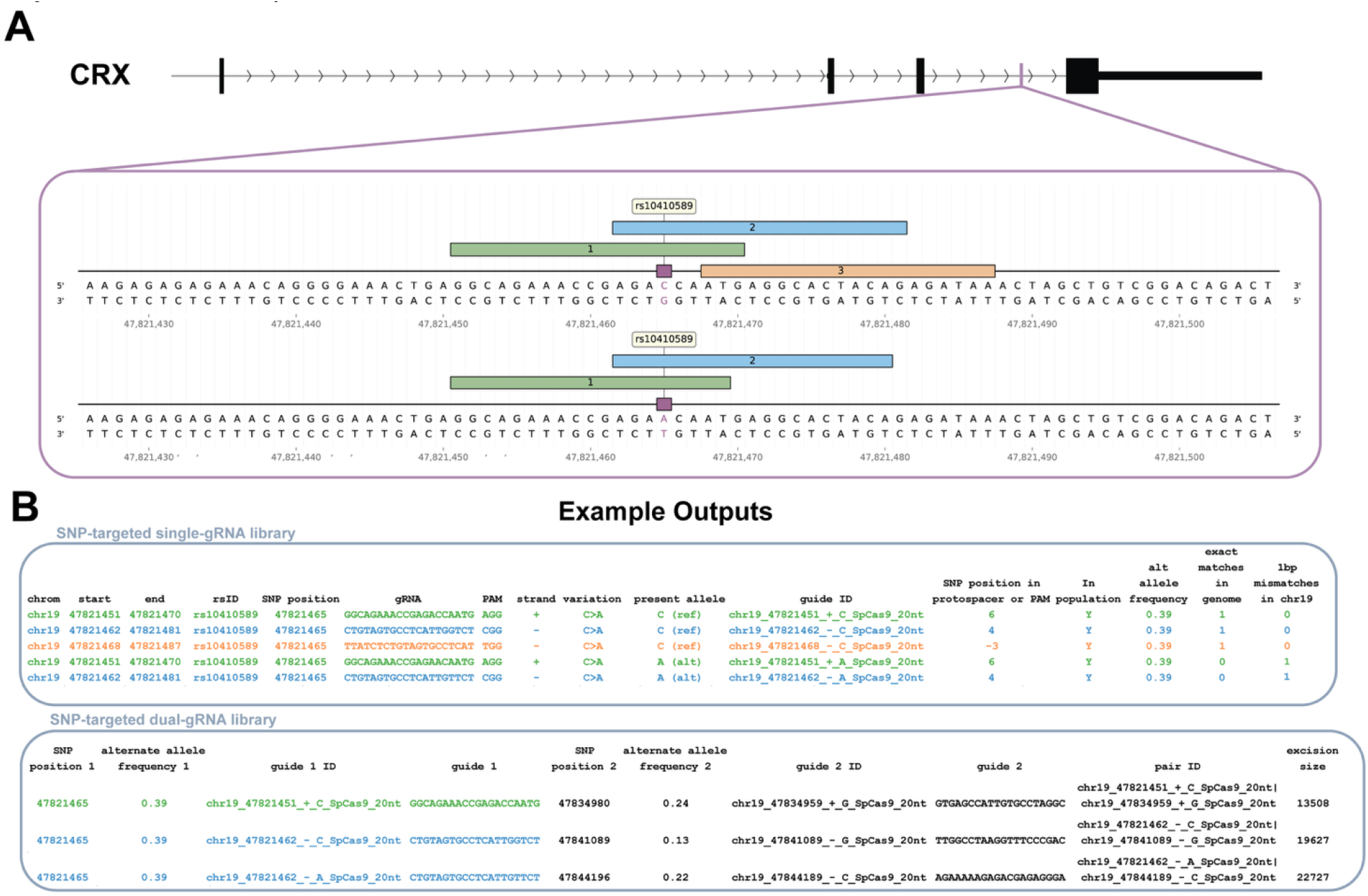
**(A)** Schematic representation of the gRNA design strategy for each allele at one region. Allele A guide sequences include the C nucleotide and allele B guides include the A nucleotide. Allele A, Guide #3 targets the ‘TGG’ PAM, which is destroyed by the SNP in Allele B. Black boxes represent gene exons. Insets: colored rectangles show gRNAs and purple rectangles represent the targeted SNP. **(B)** EXCAVATE-HT outputs a SNP-targeted single-gRNA library including SNP information, gRNA information, a unique guide ID, and off-target counting. The SNP-targeted dual-gRNA library includes SNP information for guide 1 and guide 2 as well as a unique Pair ID and excision size.

To demonstrate EXCAVATE-HT’s utility, we use it to generate libraries for the gene *CRX*, in which dominantly inherited mutant alleles cause a spectrum of retinal dystrophies. *CRX* belongs to a broader category of genes that are haplosufficient but can mutate to cause diseases with dominant inheritance patterns; we termed these genes “dominant & dispensable” or D&D^9^. For such genes, inactivating the disease-causing allele while preserving the normal allele could restore a healthy phenotype. This has already been demonstrated for dominant CRX-associated LCA7, where CRISPR/Cas9-mediated allele-specific editing was able to partially rescue photoreceptor phenotypes in patient iPSC-derived retinal organoids^10^. D&D genes are highly prevalent in neurologic, ophthalmologic, and cardiovascular diseases, motivating our selection of *CRX* as a representative use case of the EXCAVATE-HT software. Furthermore, many D&D genes harbor 100s of disease-causing missense mutations (CRX harbors 174 disease mutations), so treating each patient via mutation-targeted CRISPR editing is impractical. By enabling inactivation strategies based on common SNPs, as opposed to rare mutation-causing variants, EXCAVATE-HT would benefit large patient populations affected by this class of genetic diseases. In our recent study, we found that gene excisions have the greatest potential impact, as they can leverage non-coding SNPs to target 90% of D&D genes. In fact, many genes, such as CRX, lack common coding SNPs, so excisions are a particularly important approach. Despite the potential of this editing strategy, existing software lacks the key features for the analysis and design of allele-specific excisions. We developed EXCAVATE-HT to provide an easy-to-use tool for large-scale experimental testing of allele-specific CRISPR edits and to enable finding optimal edits by streamlining gRNA library design.

## METHODS

EXCAVATE-HT has two modes: ‘generate’ and ‘pair’. ‘Generate’ mode outputs a variant-targeted single-gRNA library for the user’s region of interest, while ‘pair’ mode takes a single-gRNA library as input and outputs a paired dual-gRNA library according to the user’s choice of pairing method. ‘Generate’ mode can also optionally pair gRNAs directly from the generated single-gRNA library, in a single, combined run. The separated ‘pair’ mode allows users to take a previously generated single-gRNA library, manually modify it as needed (e.g., remove gRNAs with many off-target effects, add positive or negative control gRNAs) before pairing.

### EXCAVATE-HT Generate: Generation of single-gRNA libraries

#### User inputs

EXCAVATE-HT Generate has the following required inputs: variant lists in variant call format (VCF) files for the individual target genome, cell line, and/or population of interest (e.g., from the 1000 Genomes Project^3^), DNA sequence in a FASTA file for the chromosome of the locus of interest, the DNA sequence for the reference genome in a FASTA file, and finally, the genomic locus of interest in the format chrom#:start-end. The user can also specify information about Cas species and PAM by either selecting one of the supported species^12^ (SpCas9, SpCas9 NG, enAsCas12a, or SaCas9) or inputting a custom set of one or more PAM sequences and their orientation. Additional optional inputs include allele frequency threshold for population-wide variant data, gRNA length, the maximum distance in base pairs between the PAM sequence and the SNP, turning on off-target counting along with accompanying inputs (see off-target counting section below for details), outputting a summary table of SNPs targeted by the output gRNA library, and splitting the gRNA library by allelic phasing. Additionally, the option to output a library of gRNA pairs can be turned on, and the pairing method can be specified (see EXCAVATE-HT Pair section below for details). User inputs go through a verification process before any analyses are run. The input Cas enzyme is initialized using saved information for the supported species, or information provided by the user about orientation and PAM sequence for a custom Cas species. This processing is done by various functions defined in ap.py.

#### Identification of SNP targets

This process is handled by the create_gens function. Variant data is loaded from input VCF files according to the type of data (individual or population). Any indel variant is excluded, and entries with missing genotypes are removed. For population VCFs, only SNPs with minor allele frequencies (MAF) above the input threshold (default 0.1) are included. For cell line VCFs, only heterozygous variants are included, and entries with missing genotypes are removed. The chromosome, position, rsID, reference and alternate alleles, and allele frequency are saved. If the cell line VCF is phased, an additional column is added to store information about which allele the gRNA targets. Variant data from each VCF input is saved in its own dataframe for the subsequent search for PAM and gRNA sequences.

#### Finding PAM and gRNA Sequences

The input chromosome file is used to read the sequence of the locus of interest. For each set of variants (each population and cell-line VCF file) that was processed as described above, an “alternate” reference genome sequence is generated by virtually mutating each SNP in the reference sequence to be its alternate form as present in the VCF file. Thus, reference and alternate sequences of the locus of interest are used to find PAM+gRNA sequences that target each respective form of the SNP. This approach ensures the capture of all allele-specific PAMs that may be created or destroyed by the different SNP alleles. The identification of PAM and gRNA sequences is done by the find_guides function. The Python regular expressions (regex) package is used to find and store the positions of all PAM sequences in the vicinity of each SNP of interest, after which only the positions that meet the SNP–PAM distance specified by the user are used. Next, the protospacer sequence adjacent to each PAM position, along with other relevant information about the gRNA, is saved in a pandas dataframe. Separate gRNA dataframes are generated for each VCF file, so they are combined, with duplicates removed and annotated with which VCF they came from by the all_guides_var_info function. If a user is interested in these separate dataframes for each VCF, they may use the –per-vcf flag to output them. If a user is interested in separating gRNA by phasing, the –split-phased flag can be used to split the combined dataframe by the phasing of the SNPs and output allele-wise libraries.

#### Off-target Counting

EXCAVATE-HT performs off-target analysis using Bowtie to identify exact genome-wide matches and 1 bp mismatches on the target chromosome. To perform off-target analysis, the user must include the --off-targets flag. Bowtie requires genome index files in .bt2 format to perform sequence alignment. There are three options to provide these index files:

1. If the user is creating libraries for the hg38 genome, they can request that pre-built hg38 Bowtie indexes be automatically downloaded and used using the --download-hg38 flag. This is the easiest and recommended option.
2. The user may provide their own existing Bowtie indexes, using the --genome-index-prefix flag followed by the full path prefix to the index files. This option is recommended when hg38 indexes need to be used for multiple subsequent runs, and have been downloaded once already.
3. If no genome index is provided or requested to be downloaded, the user must provide a genome FASTA file, and EXCAVATE-HT will build indexes for the genome automatically. This option takes 30–60 minutes, depending on the genome size.

Counts for exact matches of a gRNA in the genome and matches with up to a 1 bp mismatch in the target chromosome will be output in separate columns in the single gRNA library CSV files. Code for all off-target analysis can be found in the file off_targets.py.

#### Output files

EXCAVATE-HT Generate can produce five possible outputs:

1. A single-gRNA library, optionally split by phasing, in a comma-separated values (CSV) file. This includes information about the genomic coordinates of the gRNA, position and rsID (when applicable) of the targeted variant, the strand the gRNA targets, the alleles of the variant, the allele present in the gRNA, the PAM and gRNA sequence, a unique guide ID, the position of the variant in the protospacer, whether the variant is present in the input individual/cell line’s VCF file and/or the input population VCF file, and the variant alternate allele frequency when applicable (**Figure 2B)**.
2. Positional information about each gRNA in the single-gRNA library in the browser extensible data (BED) format. This enables input into genome browsers and viewing alongside genomic information of the user’s choice, such as ATAC-seq or ChIP-seq data, to further inform the planning or analysis of experiments using gRNA libraries generated using EXCAVATE-HT.
3. Information about each variant that is targeted in the library and a count of how many gRNAs (either reference or alternate version) target each variant in CSV format. This information is output if the --summary option is used.
4. Single-gRNA targeting variants for each input VCF in CSV format. This information is output if the --per-vcf flag is included.
5. A paired-gRNA library with unique pair IDs assigned to each pair, optionally split by phasing, along with essential information about the gRNA and targeted SNVs in CSV format (**Figure 2B)**. This file is generated only if the user opted to pair the gRNA library with the --pairing-method flag (see below).

### EXCAVATE-HT Pair: generation of dual-gRNA libraries

EXCAVATE-HT’s ‘pair’ module allows users to input a single-gRNA library and output paired-gRNA libraries.

#### User inputs and outputs

EXCAVATE-HT Pair requires as inputs a gRNA library in a CSV file and specification of the pairing method desired via the --pairing-method flag. The options for the pairing method are random (r), fixed-point (fp), or tiled (t) pairing. For random pairing, all possible pairs of gRNAs that target different SNVs are returned. Of the three, this is the most inclusive pairing strategy. In fixed-point pairing, the user must also specify one or more points in their locus of interest about which the gRNAs will be paired. The points are specified using the --fixed-points-list flag followed by one or more genomic positions without a chromosome number. This strategy is useful for the targeted excision of specific features like exons. In tiled pairing, only gRNAs that target immediately adjacent SNVs are paired, to be used in tiled screens.

## RESULTS

To identify allele-specific targeting opportunities for the *CRX* gene, we first surveyed the number of common SNPs within the coding region, including 1 kb upstream and downstream of the gene. This narrow window was chosen to avoid neighboring genes, but can be adjusted to fit the user’s needs. Here, we define common SNPs as those with a MAF greater than 10% in the 1000 Genomes Project (1kGP) cohort^3^ (**Figure 3A**).

**Figure 3.**
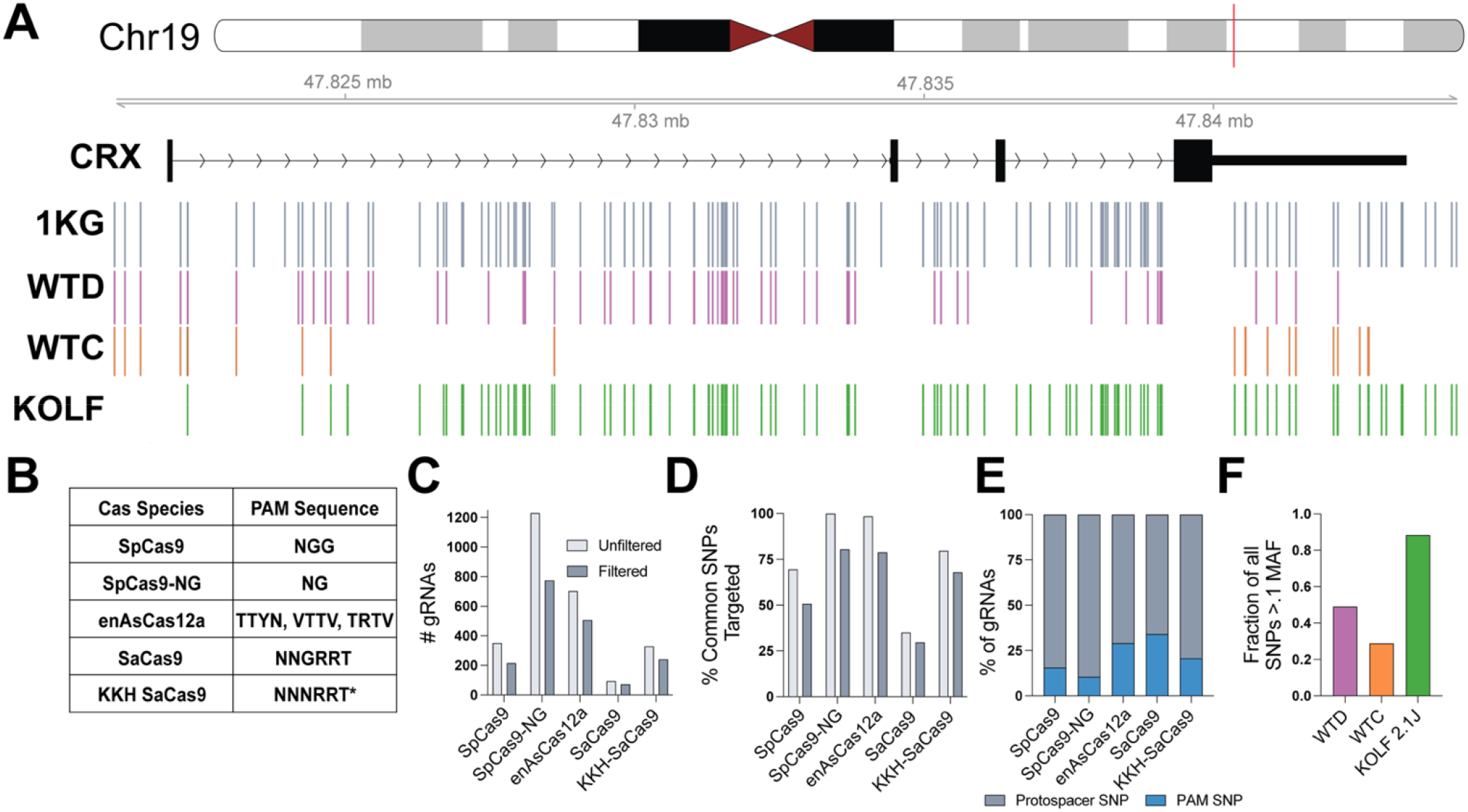
**(A)** Schematic showing chromosome 19, including the region encompassing *CRX* and 1 kb upstream and downstream. SNPs are represented as lines, colored by the cell line/population represented (grey: 1kGP Cohort, purple: WTD, orange: WTC, and green: KOLF 2.1J). **(B)** PAM sequences used to identify Cas species-specific targets,built into EXCVATE-HT Generate. * denotes custom PAM. **(C)** Number of gRNAs identified by EXCAVATE-HT before and after filtering out guides with > 1 exact genome match. **(D)** Percentage of common SNPs targeted in each Cas species specific-library before and after filtering out guides with > 1 exact genome match. **(E)** Percentage of gRNAs with SNP in PAM (blue) as compared to SNP within the protospacer. When the SNP maps to a PAM, the gRNA/Cas nuclease can only target the allele with an intact PAM. When the SNP maps to the protospacer, the PAM is intact on both alleles, and both are targeted. **(F)** Fraction of common (> 0.1 MAF) SNPs within each cell line represented in (A).

Next, we used EXCAVATE-HT Generate to identify all allele-specific SNP-targeting guides for the built-in Cas species: SpCas9, SpCas9-NG, enAsCas12a, and SaCas9 (**Figure 3B**). We also used the custom PAM feature to find gRNAs for the KKH variant of SaCas9^13^. As expected, the relaxed PAM of the SpCas9-NG variant produced the library with the greatest number of total gRNAs (1231). Additionally, this library allows for targeting 100% of the common SNPs identified. The enAsCas12a library targeted 98% of SNPs but contained 43% fewer gRNAs (**Figure 3C, D**), likely because, although the nuclease accommodates multiple PAMs, the sequences are restrictive and support few gRNAs^14^. We also found that Cas species with shorter PAM sequences offered more opportunities to design gRNAs for each SNP allele because shorter PAMs reduce the likelihood that a SNP is located within the PAM (**Figure 3E**).

When generating single-gRNA libraries, EXCAVATE-HT also outputs the number of exact matches for each gRNA in the genome as well as 1 bp mismatches on the same chromosome as the region of interest (**Figure 2B**). Our rationale for including 1 bp mismatches is that they may still allow recognition by the gRNA/Cas nuclease and, if present on the same chromosome as the target, may cause large chromosomal rearrangements. In addition, searching for off-targets is computationally laborious, and limiting the search for 1 bp off-targets to the same chromosome reduces runtime. To eliminate gRNAs with high off-target potential, we filtered out any guides with more than 1 exact match near a PAM anywhere in the genome, resulting in each library losing, on average, 30% of gRNAs but only 20% of SNP coverage (**Figure 3C, D**).

We next asked whether the single-gRNA and excision libraries generated from common SNPs in the 1kG cohort would be useful across multiple cell lines. To this end, we compared this population-level data with SNPs present in whole-genome sequenced, fully characterized wild-type cell lines: WTD^15^, WTC, and KOLF 2.1J^16^. We found that each cell line harbored >25% of the 130 SNPs present in the defined region (**Figure 3A, F**). Given that our library targets variants with a MAF greater than 10%, this level of overlap is expected and highlights the value of population-level libraries.

The filtered single-gRNA library for SpCas9 was used with EXCAVATE-HT Pair to create multiple dual-guide libraries. The random pairing method, where each guide is paired with all others, yielded over 20,000 pairs (**Figure 4A**). While this method generates the most potential pairs, fixed point pairing allows users to define excision libraries for specific regions of interest. Here, we selected the midpoint of the three coding exons (exons 2, 3, or 4), reasoning that excision of any of these exons would prevent *CRX* protein production. We avoided exon 1, because it is non-coding and its excision might not affect gene expression. This pairing method produced a library of over 14,000 gRNA pairs (**Figure 4B**).

**Figure 4.**
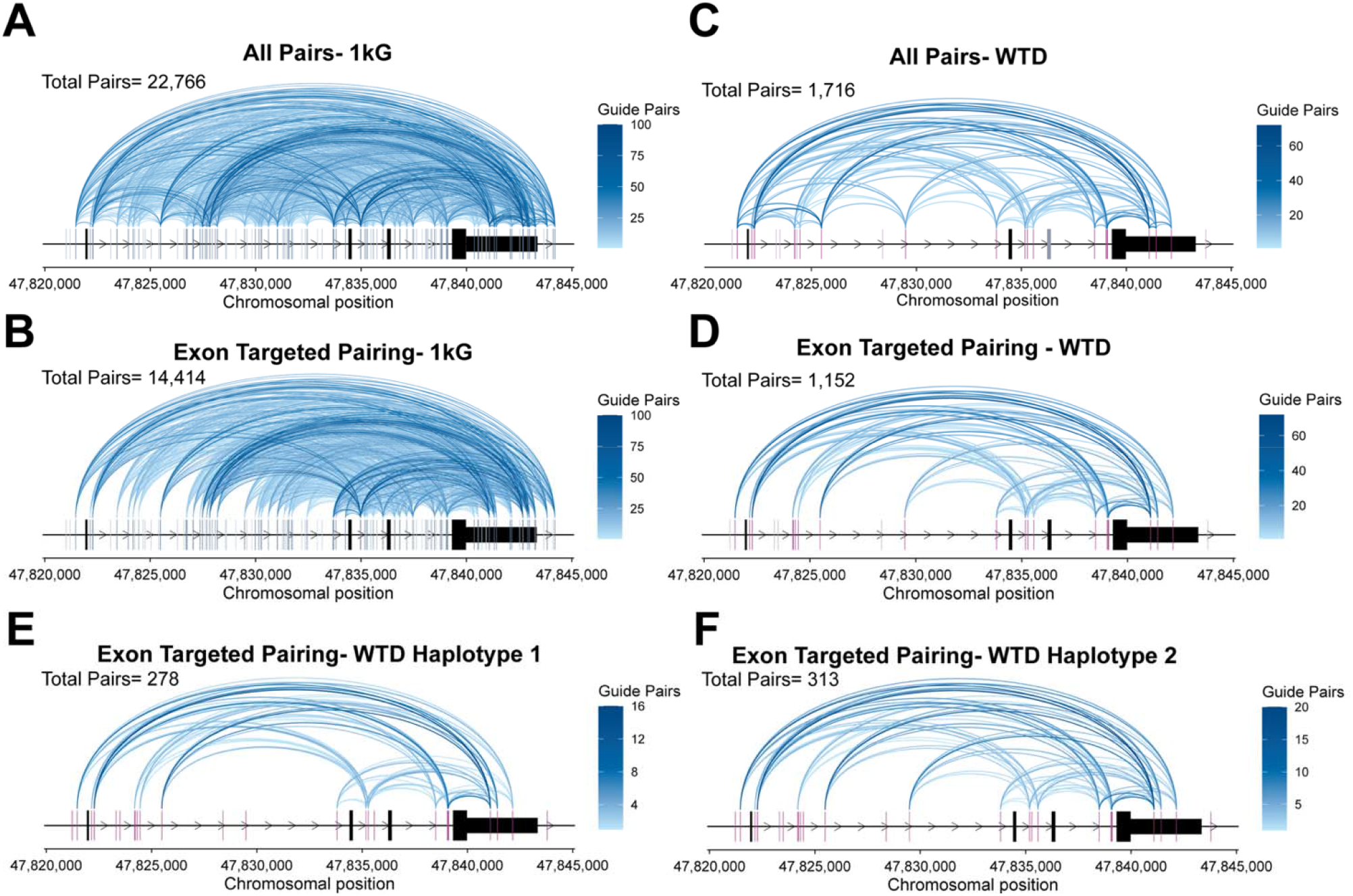
**(A, B)** Arc plots showing excision gRNA pairs generated by EXCAVATE-HT with (A) random pairing and (B) fixed point pairing methods using the SpCas9 filtered common SNP gRNA library. Dark grey lines represent targeted SNPs, light grey lines represent non-targeted SNPs. **(C, D)** Arc plots showing excision gRNA pairs generated by EXCAVATE-HT (C) random pairing and (D) fixed point pairing methods using the SpCas9 filtered common SNP gRNA library based on SNPs in the WTD cell line. Dark purple lines represent targetable common SNPs present in WTD, light purple lines represent common SNPs not targetable. **(E, F)** Arc plots showing excision gRNA pairs generated by EXCAVATE-HT with fixed point pairing methods at the A (E) and B (F) allele, using the filtered common SNP gRNA library based on SNPs in the WTD cell line separated by haplotype. For all, blue arcs represent pairs of gRNA between two SNPs, colored based on the count as defined in the color scales. Black boxes represent *CRX* exons.

While population level libraries can be utilized to validate editing strategies across multiple cell lines, EXCAVATE-HT also allows users to generate libraries based solely on cell line-specific SNPs or on cell line-specific SNPs paired with population level data. These more patient focused libraries may be utilized to eliminate noise from unproductive guides in a large pooled screen or investigate a patient-specific therapy in the case of genes with rare disease alleles. EXCAVATE-HT Generate was run with VCF inputs from the 1kGP cohort and the WTD cell line. The single-gRNA library was filtered with the same off-target threshold as above and to remove guides that targeted rare SNPs present in the cell line but not the 1kG cohort. This cell line-specific library targeting common SNPs contained 61 gRNA pairs after filtering. The single gRNA library was then used to generate excision libraries via both random and fixed-point pairing; each approach yielded a more targeted library with less potential for non-productive gRNA pairs (**Figure 4C, D**). Finally, we used phased sequencing from the WTD cell line to generate single and paired gRNA libraries split by haplotype. This results in the same number of total gRNAs, but the fixed point pairing excision libraries were smaller in size, as out-of-phase pairs are removed (**Figure 4E, F**).

Finally, to quantify the impact of haplotype editing for *CRX*, we calculated the percentage of people that could be captured by targeting common SNPs for dual nuclease editing. We first assumed that the 174 reported *CRX* mutations^11^ are inherited equally across the patient population, as no founder population has been reported.

We also assumed that SNPs are similarly distributed within the patient population as the general human population (using the 1kGP). Using the “greedy algorithm” described by Ramey et al.^9^, which prioritizes alleles targeting the most haplotypes within the population, we found that up to 77% of the population can be targeted via haplotype editing using SpCas9 (**Figure 5)**. With just 2 unique therapies (1 pair of gRNAs targeting 2 SNPs), we can reach 40% of the population. Comparing this strategy with mutation targeted editing, 2 therapies reach just 1% of patients, a 40x increase in targetability.

**Figure 5.**
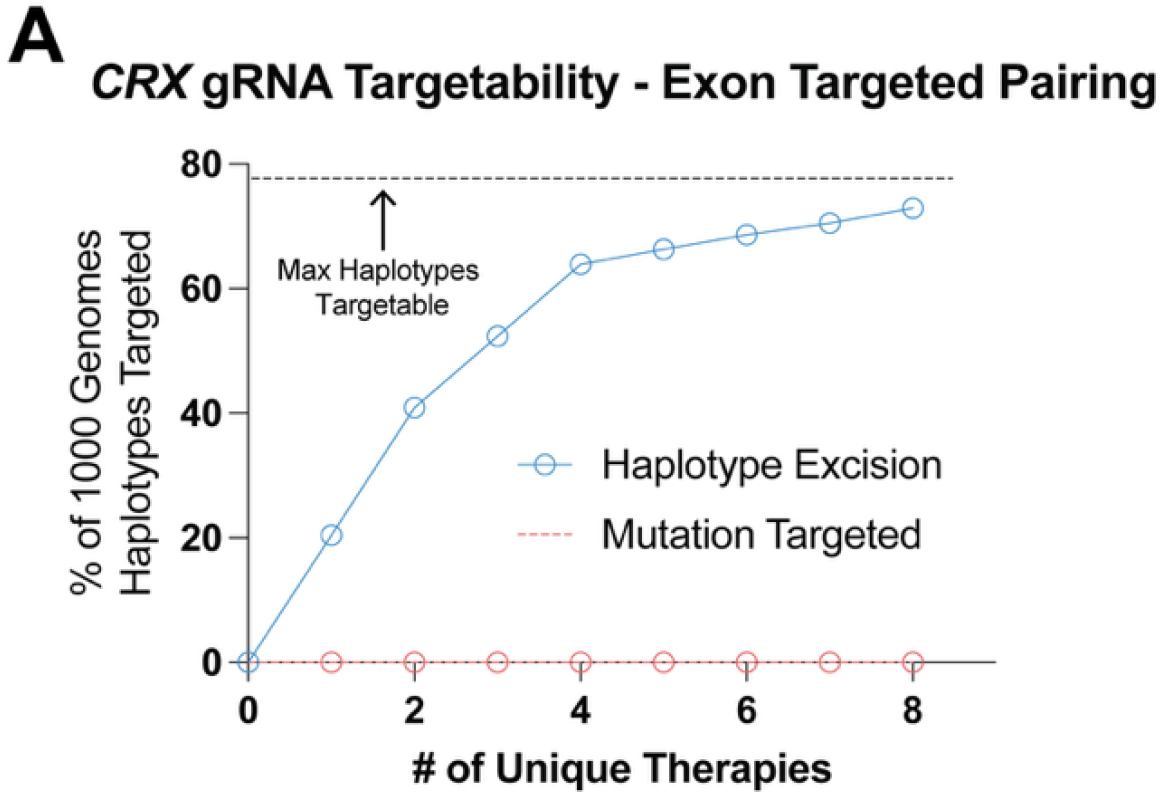
Calculation of population coverage with haplotype editing compared to mutation editing. The black dashed line represents the total cumulative target of ∼77% of 1kGP haplotypes.

## DISCUSSION

Allele-specific gene therapies targeted to common genetic variation are an untapped resource that could provide long-awaited treatments for hundreds of thousands who suffer from dominant genetic diseases. The development and testing of such haplotype-aware therapies require unbiased, large-scale screens of gRNA editing efficiency and specificity. Although this strategy holds great promise, research tools to support the design and testing of allele-specific therapeutics are lacking. EXCAVATE-HT streamlines the process of gRNA design by integrating and mining patterns of human genetic variation to spotlight high-frequency genetic variants that are targetable by CRISPR. It then uses these variants to generate a library of gRNAs that can be readily cloned, screened, and tested. Importantly, EXCAVATE-HT is easy for users to install and use locally, even those with limited bioinformatic expertise and resources.

EXCAVATE-HT’s highly flexible parameters allow researchers to design broad gRNA libraries that target all variants in a region. Recently, our group has identified over 500 genes with “dominant and dispensable” disease alleles as candidates for allele-specific CRISPR therapies. We validated the targetability of SNPs in the gene regions using the EXCAVATE-HT pipeline. We found that >90% of this gene set is targetable by allele-specific excision, which covers a much greater proportion of the patient population than the other methods of allele-specific editing (epigenetic silencing, exon disruption, and splice disruption). For all genes in this test set, excision offers the greatest opportunities for allele-specific targeting because intergenic regions harbor a higher frequency of SNPs. Identification of the most valuable of the 1000’s of SNP-enabled excisions by hand is tedious, and researchers may miss opportunities by only selecting a handful. EXCAVATE-HT enables researchers to computationally design gRNAs easily, without the need for genome browsing software. While this workflow is beneficial for designing therapeutic excision gRNAs, EXCAVATE-HT can also be used to identify excision gRNAs for generating heterozygous knockout cell lines for disease models, a valuable resource in biomedical research.

In addition to targeting all possible SNPs, EXCAVTE-HT can be used to target common SNPs, enabling a single gRNA library (single or excision) to work both therapeutically across the patient population and experimentally across multiple cell lines. The value of targeting common SNPs is apparent in the CRX gene, which harbors >170 disease mutations. We tested this using 3 wild-type cell lines as compared with SNPs from the 1kGP. Every cell line harbored a subset of these SNPs, demonstrating that using libraries generated from EXCAVTE-HT could be tested across all three without the need to design multiple libraries for each. That said, because EXCAVATE-HT allows users to generate libraries from multiple VCF files (populations and multiple cell lines) simultaneously and to export an editable gRNA library, users can filter the output to create unique, cell-line-specific libraries if desired.

In this example, using the EXCAVTE-HT software, we generated multiple excision gRNA libraries to disrupt a single allele of the disease-causing gene *CRX*. This library can be used directly to conduct an unbiased screen for guide efficiency and specificity. The rules governing large, dual-guide excisions have yet to be explored, with potential contributing factors including cell type, guide sequence, PAM orientation, linear distance, and 3D genomic distance. EXCAVATE-HT is well-suited to empower screens to answer these open questions.

To summarize, EXCAVATE-HT adds to a landscape of gRNA generation tools that support the discovery of efficient and specific CRISPR-based edits. EXCAVATE-HT enables the screening and development of allele-specific CRISPR therapies that leverage commonly inherited genetic variation, aiming to democratize and facilitate large-scale editing experiments for novel therapeutic discovery.

## ACKNOWLEDGEMENTS

We would like to thank all members of B.R.C’s & J.A.C.’s labs for helpful feedback. G.D.R. was supported by NIH grant F31AG090013 and the UCSF Discovery Fellows. J.A.C. was supported by NIH R35GM127087 and CIRM DISC0–17363. B.R.C. received funding from CIRM INFR6.2–15527 and CIRM DISC0–17363. B.R.C. acknowledges generous support through gifts from the Roddenberry Foundation and Pauline and Thomas Tusher. B.L.M. received support from CIRM EDUC4–12766G, NIH grant F32AG081085, and L’Oréal USA for Women in Science Program. We also thank Francoise Chanut for valuable editorial and scientific insights. The contents of this publication are solely the responsibility of the authors and do not necessarily represent the official views of NIH, CIRM, or any agency of the US government or the State of California.

## AUTHOR CONTRIBUTIONS

AGS, GDR, JAC, BRC, and BLM conceived the project. AGS developed the software with assistance from GDR, under the supervision of BLM, BRC, and JAC. BLM and AGS wrote the manuscript and created visualizations with revisions from all authors.

## CODE AVAILABILITY

All scripts necessary to execute EXCAVATE-HT workflows are located at https://github.com/akshitasax/EXCAVATE-HT^16^, with detailed installation and running instructions at https://excavate-ht.readthedocs.io/en/latest/.

